# UDIST: unsupervised disentanglement of shape and texture for multi-scale phenotypic profiling in 2D microscopy

**DOI:** 10.64898/2026.07.26.740036

**Authors:** Bram M. Bosch, Maarten L. Terpstra, Matthew B. Smith, Krijn H. van der Steen, Caspar T.H. Jonker, Vitalijs Ovcinnikovs, Thomas H. Wesselink, Anne F.J. Janssen, Lianne Winkel, Eva M. A. Huigen, Juliet W. Lefferts, Enrico Mastrobattista, Edo D. Elstak, Cornelis A.T. van den Berg, Jeffrey M. Beekman, Sam F. B. van Beuningen

## Abstract

Microscopy-based phenotypic profiling relies increasingly on autonomous, unsupervised feature extraction, yet no existing method explicitly separates shape from texture into dedicated and independent latent subspaces by architectural design. Therefore texture, encoding critical biological information such as protein distribution and intracellular organisation, remains inaccessible as an independent feature domain in standard unsupervised approaches. This represents a fundamental limitation that prevents unbiased phenotypic analysis across biological scales. Here we introduce UDIST (Unsupervised Disentanglement of Shape and Texture), a sequential dual variational autoencoder (VAE) framework that tackles this fundamental limitation by explicitly decoupling shape from texture into independent, non-overlapping latent subspaces at the single-object level. By training two VICReg-regularised VAEs on principal-axis-aligned objects, UDIST separates binary shape from continuous texture information into rotation-invariant feature spaces, enabling separate downstream analysis of both domains. We validated UDIST across biological scales, from nuclei and single cells to patient-derived intestinal organoids, using both fluorescence and brightfield imaging, revealing phenotypic differences previously hidden by morphological variation and enabling the independent analysis of shape and texture in downstream analyses including clustering and similarity measurements. UDIST provides a versatile, label-free, and unsupervised tool for multi-scale phenotypic profiling in high-content microscopy and screening.

## Introduction

Microscopy based phenotypic assays enable quantitative analysis of cellular states across low and high-throughput experimental set-ups. Traditionally, this depends on predefined and often handcrafted computational features, such as cell and organelle - size, -shape, and -intensity to determine phenotypic differences between experimental conditions. Tools like CellProfiler support researchers to analyse these features for microscopy images in high throughput analysis pipelines (Carpenter et al., 2006). However, to capture the full complexity of heterogeneous cell systems, the field is increasingly making use of Artificial Intelligence (AI) and deep learning frameworks. (Burgess et al., 2024; Moshkov et al., 2024). Rather than relying on predefined features, these methods autonomously learn complex feature sets directly from the raw data, enabling detection of subtle and complex morphological patterns and phenotypic relationships beyond what can be manually defined.

Among these deep-learning framework, the Variational Auto-Encoder (VAE) is well established for its autonomous unsupervised feature extraction in both general-purpose and biological image analysis (Kingma & Welling, 2013; Lafarge et al., 2019). By learning a compressed latent space representation through self-supervision for each image, these VAE’s map the raw pixel data into a compact yet descriptive set of features for any given dataset (Kingma & Welling, 2013). Analysing the relative positions of these vectors within the latent space allows researchers to get better insights into biologically meaningful relationships between samples.

An unresolved challenge of these unsupervised mapping approaches for biological image analysis is the explicit separation of cellular shape from texture features. Although both these distinct feature domains inherently co-exist within all microscopy images, no current method explicitly separates them into dedicated and independent subspaces. While recent work has explored emergent disentanglement of morphological attributes including shape and texture within a shared latent space (Murthy et al., 2025), explicit architectural separation into guaranteed independent subspaces remains unaddressed. Shape and texture frequently correlate in biological samples - for example, when intensity patterns follow cellular boundaries. However, they differ fundamentally in biological origin. Shape encodes cellular morphology and boundary constraints, whereas texture captures intracellular variations in continuous signal patterns, like protein density and distribution. Without the separation of these phenotypic feature domains, dominant variations in cell shape can heavily confound these unsupervised image analysis methods, thereby potentially masking subtle yet meaningful biological information such as a distribution change in intensity patterns.

Although standard VAE architectures have pushed the boundaries of automated feature extraction, they fundamentally struggle with true feature disentanglement: the capacity to encode isolated features into distinct, independent dimensions of the latent space. In addition, previous work showed that for both explainability and analysis, one needs strong feature disentanglement in the latent space (Adel et al., 2018; Bengio et al., 2012). Improving the latent space structure, and to some extent disentanglement, typically requires the explicit enforcement of regularizations or structural constraints (Bardes et al., 2021; Higgins et al., 2016). However, true unsupervised disentanglement without inductive biases has been shown to be theoretically impossible (Locatello et al., 2019). For biological image analysis this limitation implies that standard architectures cannot autonomously decouple cellular morphology (shape) from internal cellular signal patterns (texture). An effective solution therefore requires constructing explicit separate subspaces of shape and texture within the latent space, achieving disentanglement by architectural design rather than by constraints alone. Such separated feature spaces would enable independent analysis of both shape and texture in downstream tasks such as clustering or classification.

Here, we introduce UDIST (Unsupervised Disentanglement of Shape and Texture), a novel autoencoder training framework specifically designed for the disentanglement and separation of shape and texture features. We demonstrate that creating distinct subspaces for shape and texture enhances downstream analysis across information domains when compared to conventional VAE methods. We further introduce a novel feature-cleaning method using Principal Component Regression that utilises the partitioned latent spaces. Using a custom synthetic dataset containing three shape classes and three non-shape-correlated texture classes, we show that UDIST substantially improves downstream analytical performance.

To establish biological relevance and multi-scale adaptability, we perform a comprehensive analysis across three datasets spanning distinct biological scales, nuclei, cells and organoids, as well as two imaging modalities, fluorescence and brightfield. By using standard downstream analyses including clustering, similarity measurements and classification, we demonstrate that the UDIST framework generalises effectively across diverse biological contexts.

## Results

### Shape dominates texture in standard unsupervised feature extraction

Microscopy images of cells intrinsically contain both shape morphology and internal cellular signal patterns or texture (fluorescence or transmitted light). When analysing these images in an unsupervised manner, we observe that standard single object feature extraction methods struggle with respecting both domains and often get dominated by shape information. This is in contrast with natural image learning where it has been often observed that texture is the dominating feature (Baker et al., 2018; Geirhos et al., 2018; Hermann et al., 2020). To demonstrate this problem in a controlled setting we have created a microscopy-like synthetic dataset with three distinct shape classes (round, elongated, or protrusive) and three distinct texture classes (empty, sparse puncta, or dense puncta) which are crossed independently to create all 9 combinations (Figure 1A). We used a standard VAE architecture with 256 latent dimensions to perform feature extraction on the synthetic dataset. In the PaCMAP based dimensionality reduction plot, the dense puncta texture class can be separated from other texture classes while respecting shape information (Figure 1B). However, empty and sparse puncta texture patterns cannot be separated from each other and from different shape patterns. We measure the distances in the 256 dimensional latent space using the Sliced Wasserstein Distance (SWD) over 256 slices to quantify our observations (Flamary et al., 2021; Rabin et al., 2011). This shows the same trend: the dense texture class is far removed from both empty and sparse texture classes. Additionally, the SWD plot shows barely any separation between the sparse and empty texture class samples, instead focusing on the underlying shape classes. This indicates that standard VAEs used for unsupervised feature extraction, can separate shape patterns and dominant texture patterns, but are failing with more subtle texture patterns. In microscopy images of cells, different objects of interest can often be identified by these descriptive yet subtle texture patterns (e.g. fluorescent signal distribution). Moreover, shape features can act as independent variables due to the nature of cell growth and shape. All together, these observations show a gap in existing feature extraction methods for image datasets where shape acts as an independent variable (Figure 1C). Therefore, we propose a model architecture that promotes the separation of the binary shape information from continuous texture information into separate subspaces – for shape and texture – by explicit model design (Figure 1D). This produces cleaner, domain-specific feature representations, eliminating the need for implicit negotiation by the model between shape and texture features. It thereby allows for independent analysis of both shape and texture without loss of biologically relevant information.

**Figure 1:**
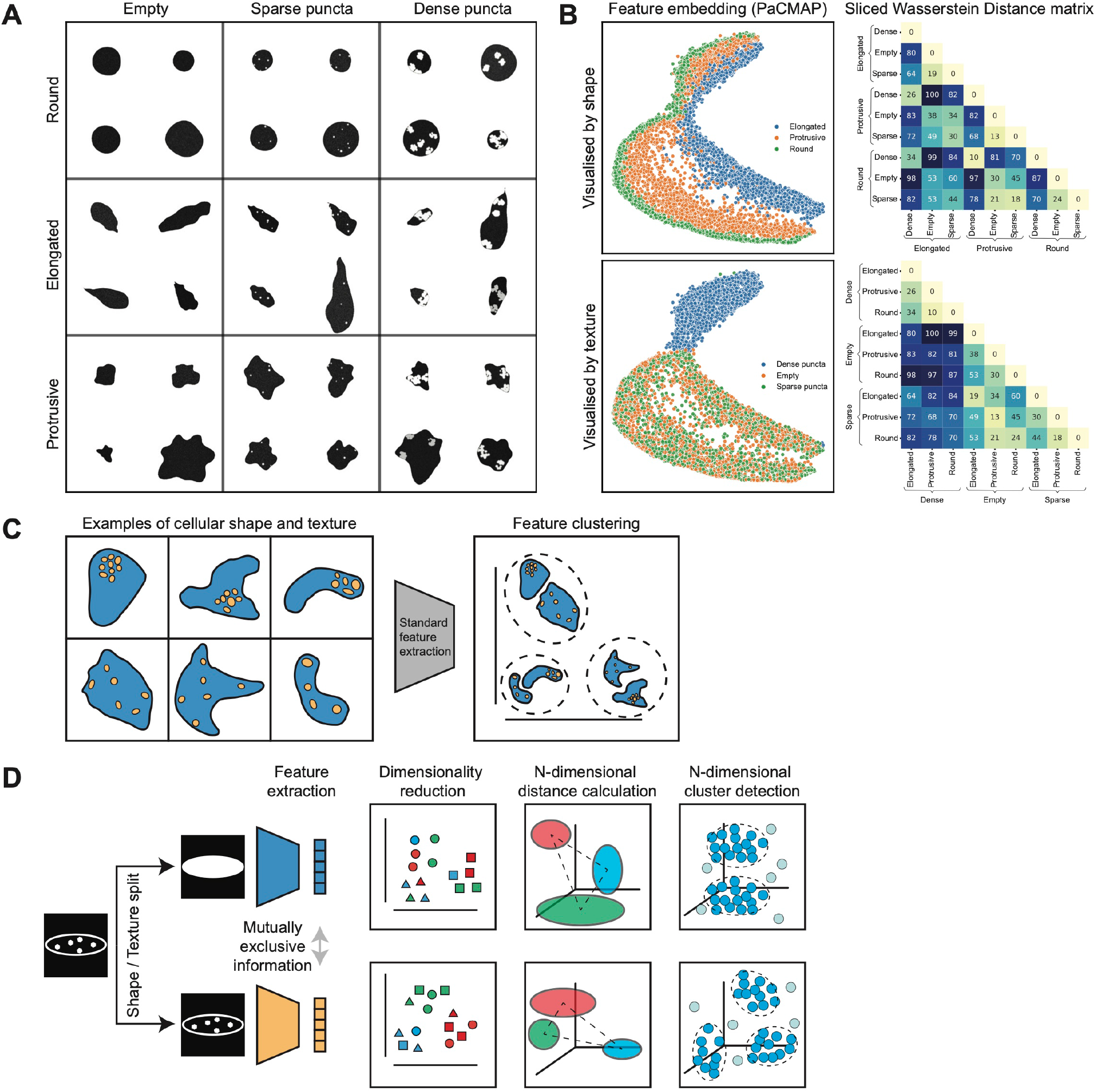
Shape dominates texture in standard VAE-based feature extraction. **(A)** Example objects from the synthetic dataset, comprising three shape classes (round, elongated, and protrusive) and three texture classes (empty, sparse puncta, and dense puncta) combined across all nine possible combinations. **(B)** PaCMAP embedding of the latent space generated by a standard VAE, coloured by shape label (top) and texture label (bottom), where each point represents a single synthetic cell. Inter-cluster distances were quantified using the Sliced Wasserstein Distance. **(C)** Schematic illustration of the shape-texture entanglement problem in standard unsupervised feature extraction for microscopy images. **(D)** Schematic of the proposed solution, in which shape and texture are explicitly separated into independent latent subspaces by architectural design.

### UDIST achieves shape-texture disentanglement through two-step VAE training

To overcome the demonstrated challenge of entanglement of shape and texture features in microscopy images using standard unsupervised feature extraction methods, we introduce a model architecture specifically designed for Unsupervised Disentanglement of Shape and Texture (UDIST). UDIST is a novel two step training approach using VAEs as basis for unsupervised feature extraction of both shape and texture independently in 2D biological images (Figure 2A). First, single cells are segmented, and the resulting binary masks are used to train a shape-based VAE model. By first training this shape-based VAE, the model learns a comprehensive and representative latent space for the shapes in the presented dataset. Secondly, the weights of the trained shape model are frozen, and a second VAE is trained on single cell texture information using the segmented masks as crop on the original texture channel. During this second training we additionally encode the masks again using the shape model. Using variance, invariance and covariance regularisation (VICReg) (Bardes et al., 2021) the new texture latent space is discouraged to encode any information already present in the shape latent space. The decoder from the new model is fed both the new texture encoding and the pretrained shape features. After training the extracted features from the trained shape encoder and trained texture encoder are used during inference for further analysis.

**Figure 2:**
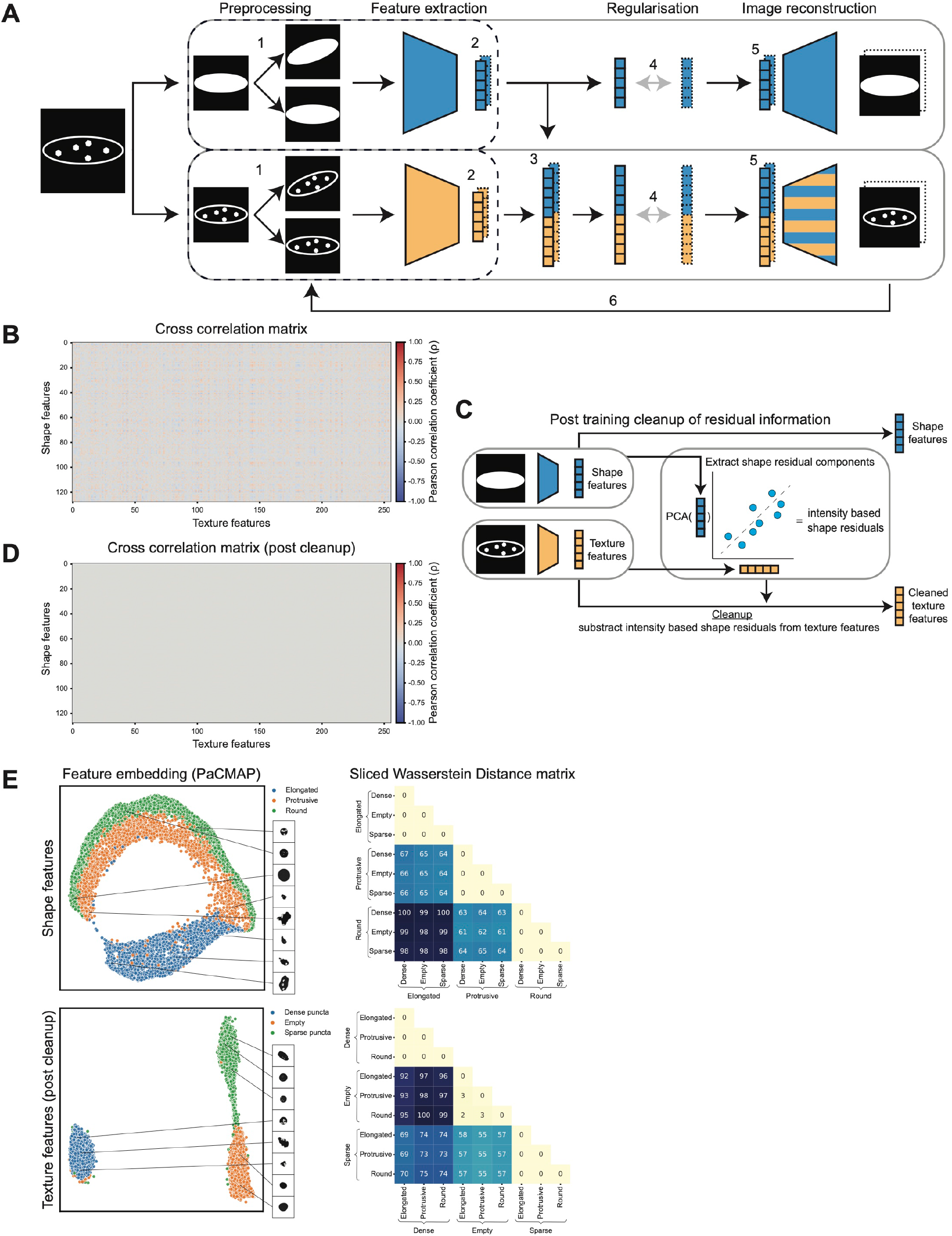
UDIST explicitly separates shape and texture into independent latent subspaces. **(A)** Schematic overview of the UDIST training procedure. (1) The input image is randomly rotated between 0 and 20 degrees. (2) Shape and texture features are extracted using independent encoders. (3) Shape and texture features are concatenated. (4) Inter-subspace independence is encouraged via VICReg regularisation (variance, invariance, and covariance terms). (5) The original image is reconstructed using a decoder. (6) Reconstruction quality is assessed using MAE and MS-SSIM. **(B)** Schematic overview of the post-training Principal Component Regression-based cleanup step for removing residual shape information from the texture latent space. **(C)** Cross-correlation matrices showing Pearson correlation coefficients between shape and texture latent dimensions for the synthetic dataset, before and after the cleanup step. **(D)** PaCMAP embedding of UDIST shape (top) and post-cleanup texture (bottom) latent spaces for the synthetic dataset, where each point represents a single synthetic cell. Inter-cluster distances were quantified using the Sliced Wasserstein Distance.

However, due to the nature of shape and texture information being inherently derived from the same object and the model having to balance a combination of sometimes competing loss functions. We expect and observe some leakage of shape information into the texture latent space (Figure 2B). Therefore, to further optimize disentanglement between the subspaces, we implement Principal Component Regression to identify and clean any residual shape information from the texture latent space (Figure 2C). After this cleaning step, any linear correlation between the latent dimensions of the separate subspaces is completely removed (Figure 2D). To validate our approach, we trained a UDIST model with 128 shape dimensions and 256 texture dimensions on 28,350 cells from the synthetic dataset described earlier. Next, we analysed the shape and texture feature latent spaces separately again using the PaCMAPs and Sliced Wasserstein Distance methods on a test set of 8,100 cells. The shape features show a clear independence from texture features (Figure 2E), as is expected since the shape features were generated from binary masks that exclude any texture information. Furthermore, the post-cleanup texture features show a specific clustering of texture classes independent of shape. Altogether, this indicates that UDIST can successfully disentangle texture features from shape features allowing for the separate analysis of these information domains.

### UDIST resolves EGFR endosomal fate from EGFR-GFP signal distribution alone

To demonstrate the utility of UDIST in a biologically relevant context we analysed imaging data from a screen employing therapeutic antibodies that may induce endocytosis of EGFR on EGFR-GFP expressing A549 cells (Lieber et al., 1976). Endosomal pathways have traditionally been characterized using co-staining approaches (Ding et al., 2006; Laniel et al., 2025; Roepstorff et al., 2009; van der Beek et al., 2021). However, previous work has shown that the spatial distribution of endosomes within the cell is itself indicative for the predominant endosomal pathway engaged (Roepstorff et al., 2009; Sorkin & Goh, 2009). We therefore hypothesised that the intracellular distribution of EGFR-GFP signal, captured as a texture feature, could serve as a read out for the dominant endosomal pathway under different treatment conditions, without the need for additional co-staining.

To test this hypothesis, we trained a UDIST model on a training set of 82,551 individual cells spanning a range of conditions, followed by texture feature extraction on a test set of 30,702 individual cells encompassing 6 experimental and 3 control conditions. The first control condition comprised of untreated cells, representing a steady state distribution of EGFR-GFP. The second control condition are cells imaged one hour after EGF stimulation, representing an early endosomal state (Sorkin & Goh, 2009). The third control condition contain cells imaged four hours after EGF stimulation, representing a late endosomal state (Sorkin & Goh, 2009). The post-cleanup texture latent representations were then used to characterise the EGFR-GFP signal distribution across control conditions. As shown in Figure 3A, individual cells form condition specific clusters in PaCMAP space, with each control conditions occupying a distinct region of the embedding. Partial overlap between conditions was also observed, reflecting the inherent biological heterogeneity of EGFR-GFP distribution within individual conditions. Notably, shape features showed a uniform, non-specific distribution across conditions (Figure S1A-B), indicating that cell morphology is not systematically influenced by experimental treatments.

**Figure 3:**
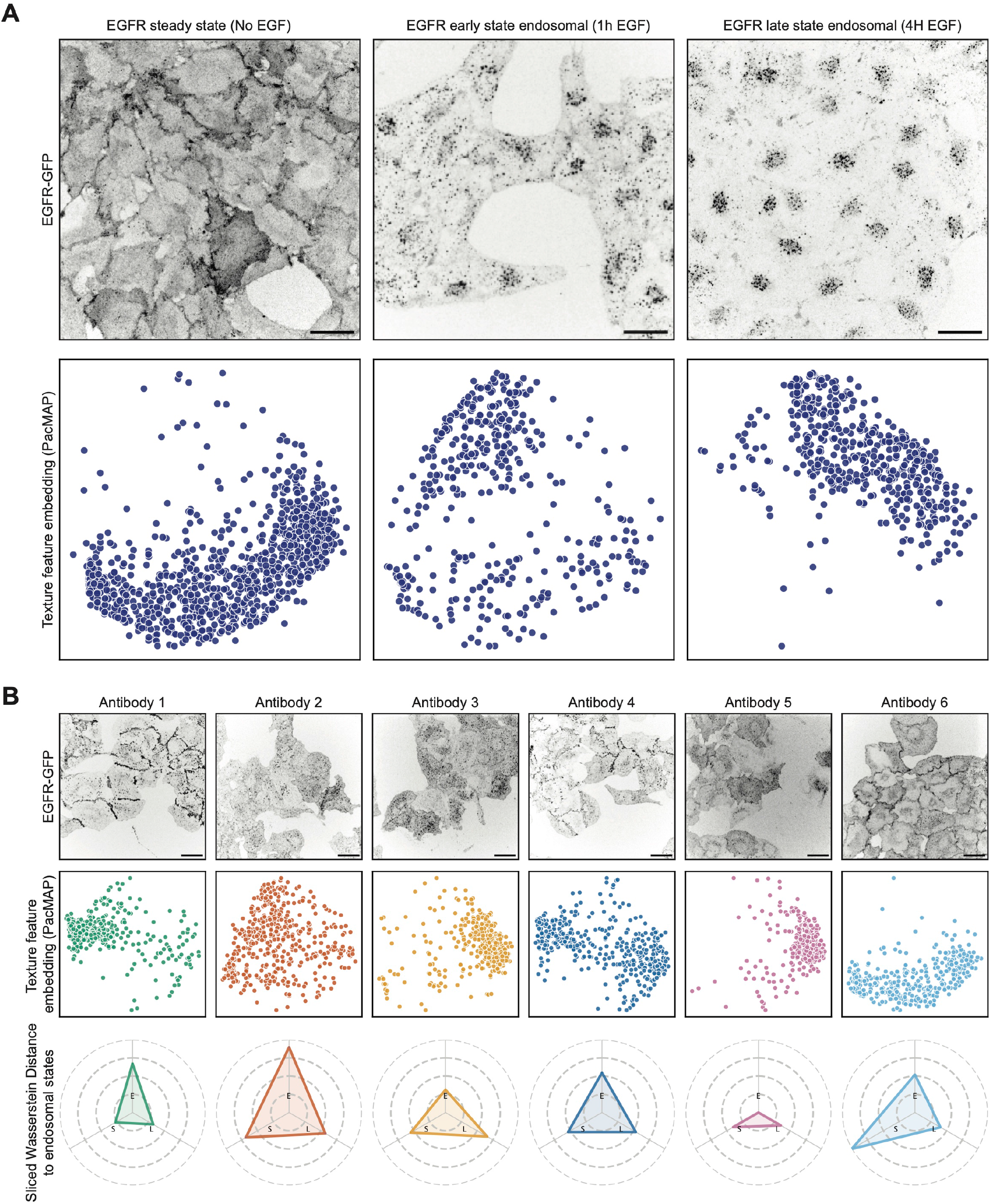
Marker-independent endosomal pathway analysis in A549 cells using UDIST. **(A)** Representative examples of A549 cells expressing EGFR-GFP for each control condition (upper pannel) and PaCMAP embedding of post-cleanup texture latent features for all cells in the control conditions, where every point represents a single cell (lower panel). **(B)** Representative examples of A549 cells treated with distinct antibodies targeting EGFR-GFP (upper panel), PaCMAP embedding of the post-cleanup texture latent features where every point represents a single cell (middle panel), and similarity scores of each experimental condition relative to each control condition, calculated in the original latent space using the min-max Sliced Wasserstein Distance (lower panel). S: steady state, E: early endosomal, L: late endosomal. Scale bars = 30 um. All microscopy images are displayed as inverted intensity for clarity. PaCMAP embeddings show a random subsample of 10% of all cells.

To assess phenotypic differences, we compared these control conditions of known EGFR-GFP distribution phenotype against experimental conditions of unknown phenotype induced by distinct anti-EGFR antibodies (Figure 3B). To quantify these phenotypic differences, we employed the min-max implementation of the Sliced Wasserstein Distance, whereby the texture feature similarity of each experimental condition is calculated relative to each of the three control conditions, represented along three axes (S: steady state, E: early endosomal, L: late endosomal). Antibody 1 shows high membrane localization and clear puncta in the image. The analysis shows a clear similarity to E, indicating slow but significant internalization that is not reaching late compartments during the course of the experiment. Antibody 2 shows no membrane staining and a diffuse punctate pattern in the image. The analysis shows a good similarity with E, L and S, indicating an efficient uptake from the membrane without preferential sorting to the degradative pathway. Antibody 3 shows no membrane staining and a diffuse punctate pattern similar to antibody 2 in the image. However, the analysis shows more similarity towards L, indicating that in contrast to antibody 2, here a preferential sorting to the degradative pathway is present. Antibody 4 shows high membrane localization and clear puncta in the image, similar to antibody 1. However, a higher similarity to L here indicates late endosomal compartments are reached more efficiently. Antibody 5 treated cells exhibit a phenotype distinct from all three controls, suggesting engagement of an endosomal pathway not represented within the current control set. Finally, antibody 6 treated cells show a high similarity to the steady state control, consistent with normal EGFR-GFP trafficking. Collectively, these results demonstrate that the prominence of distinct pathways can be inferred solely from the intracellular EGFR-GFP signal distribution, without the need for pathway specific co-stainings. Importantly, computing similarity against all three controls simultaneously, rather than assigning a single categorical label, preserved the phenotypic heterogeneity inherent to the samples. Conventional endosomal pathway assays typically rely on exogenous markers such as Dextran and Transferrin to demarcate early, late and recycling endosomal compartments. Our results demonstrate that UDIST can circumvent this requirement by capturing and quantifying the visual phenotypic similarity of the single cell EGFR-GFP signal directly. This offers a streamlined and marker independent approach to endosomal pathway analysis.

### Unsupervised clustering of PML-II nuclear condensate phenotypes in nuclei using UDIST

Promyelocytic leukemia protein (PML) has been shown to form condensates on the nuclear envelope during lamina disruption (Janssen et al., 2026; Jul-Larsen et al., 2010; Ohsaki et al., 2016). While PML isoform I predominantly gives rise to punctate condensates, PML isoform II (PML-II) forms a variety of nuclear condensate phenotypes. Characterising these phenotypic subtypes is complicated by the heterogeneous nature of the signal and by inter-nuclear variation in shape, which can confound the detection of subtle differences in condensate patterns. As UDIST explicitly separated shape from texture, it should isolate condensate pattern variation from confounding morphological differences between individual nuclei.

To exploit this capability, we applied UDIST to a microscopy dataset of GFP-PML-II overexpressing HeLa cells to cluster condensate pattern subtypes in an unsupervised, annotation-free manner. The dataset comprised a control condition and five knockdown conditions, in which individual proteins of interest – identified from a BioID screen for PML-II interactors (Roux et al., 2018) – were depleted in these HeLa cells using siRNAs (Figure 4A). In total 71,642 nuclei were used for model training and 2,034 nuclei for inference. Given the exclusive focus on condensate texture patterns, shape features were discarded and only post-cleanup texture representations were retained for downstream analysis. Consistent with this, PaCMAP visualisation of the shape feature space confirmed a uniform distribution across knockdown conditions, indicating that nuclear shape is not systematically influenced by the experimental perturbations (Figure S2). In contrast, the texture space revealed that controls (siCTRL) were broadly distributed across the projection, consistent with the phenotypic heterogeneity characteristic of PML-II condensates (Figure 4B). Furthermore, each knockdown condition occupied a distinct subregion of the embedding, indicating that depletion of specific interactors systematically shifts the prevalence of particular condensate phenotypes. To identify discrete condensate pattern subtypes, Leiden clustering, a community detection algorithm (Traag et al., 2019), was applied to the texture latent space, yielding six clusters (Figure 4C). Each cluster captured a distinct phenotype: cluster 0 comprised of nuclei with a dense, amorphous condensate pattern; cluster 1, nuclei densely populated with discrete puncta; cluster 2, nuclei with sparse puncta; cluster 3, nuclei with absent or very low signal; cluster 4, a heterogenous group enriched for outliers from all other clusters; and cluster 5, nuclei characterised by one or more intensely bright patches (Figure 4D). Quantification of cluster enrichment relative to siCTRL revealed distinct over- and underrepresentation of condensate pattern subtypes across knockdown conditions (Figure 4E). Knockdowns 1 and 4 shared a similar pattern: both were underrepresented in cluster 0 (chaotic condensate patterns) and overrepresented in cluster 2 (sparse puncta). In contrast, knockdowns 2 and 3 showed the inverse relationship, being overrepresented in the chaotic condensate cluster (cluster 0) and underrepresented in the sparse puncta cluster (cluster 2), as well as overrepresented in the bright patch cluster (cluster 5). Finaly, knockdown 5 resembled knockdowns 1 and 4 in its overrepresentation in the sparse puncta but did not show underrepresentation in the chaotic condensate cluster, suggesting a partially distinct mechanism of action.

**Figure 4:**
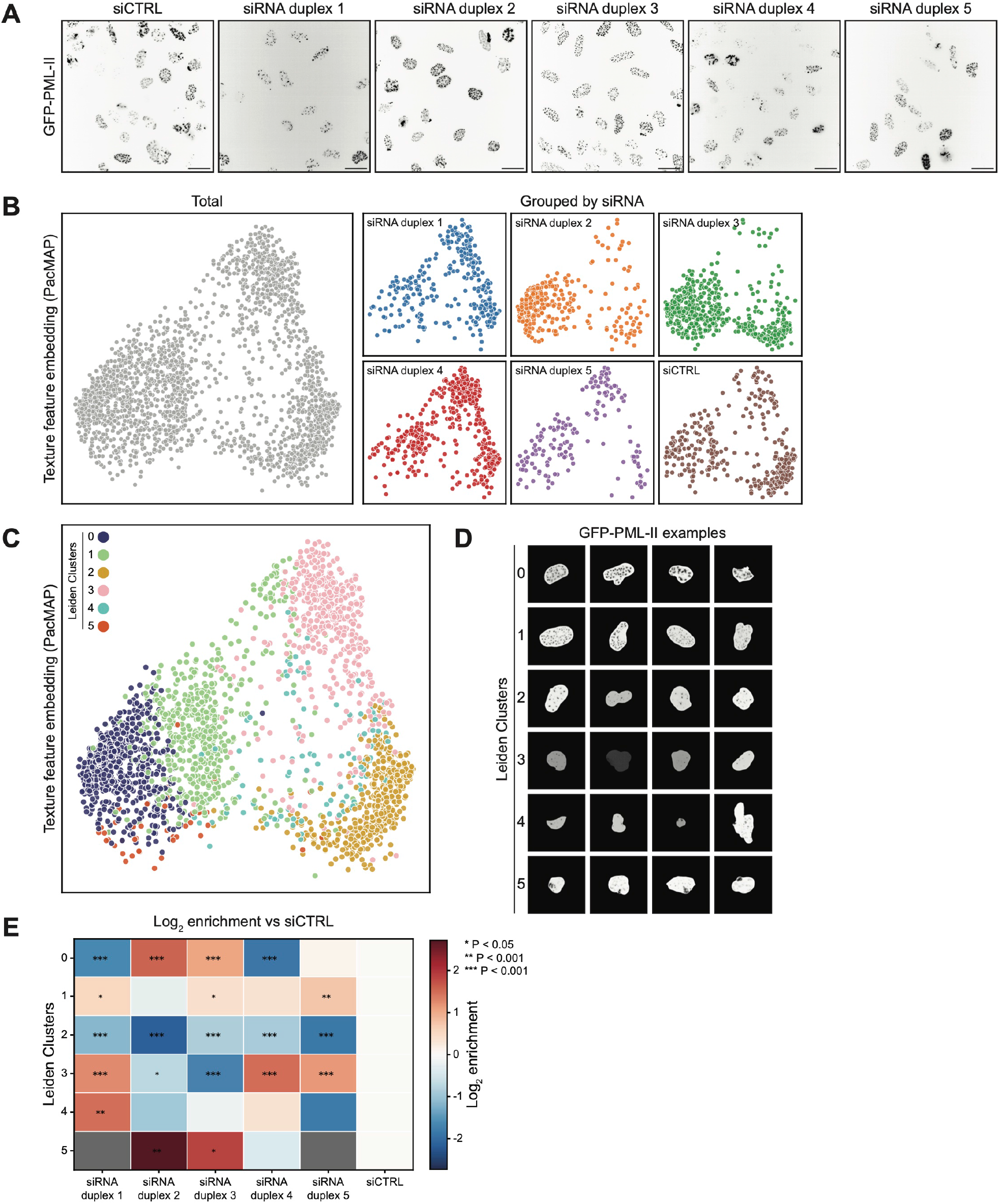
Unsupervised clustering of GFP-PML-II nuclear condensate phenotypes in HeLa cells. **(A)** Representative examples of HeLa cells expressing GFP-PML-II for the siCTRL condition and each of the five siRNA conditions. **(B)** PaCMAP embedding of post-cleanup texture latent features where every point represents a single nucleus (left panel), and PaCMAP embeddings showing the localisation of each siRNA condition within the texture latent (right panels). **(C)** PaCMAP embedding of all nuclei labelled by Leiden clusters assignment. **(D)** Representative example nuclei for each detected Leiden cluster. **(E)** Quantification of the distribution of siRNA condition across Leiden clusters, normalised to the siCTRL. Grey panels indicate Leiden clusters containing no nuclei from that condition.. Scale bars = 30 um. All microscopy images are displayed as inverted intensity for clarity.

Collectively, these results demonstrate that VAE can capture and cluster nuclear intensity pattern subtypes in an unsupervised, annotation free, and shape independent manner. Thereby enabling the discrimination of biologically meaningful condensate phenotypes across experimental conditions.

### Brightfield based phenotypic profiling of wildtype and cystic fibrosis organoids using UDIST

As a final experimental dataset, UDIST was trained and evaluated on brightfield images of patient derived intestinal organoids (PDIOs). The PDIOs were obtained from donors with either wild-type *CFTR* or disease-causing *CFTR* variants, with all Cystic Fibrosis (CF) samples derived from patients homozygous for the F508del mutation (F508del/F508del). Prior work has established that organoid morphology is defined by CFTR-dependent luminal ion and fluid secretion, and discriminates between healthy and CF and CFTR modulating treatments (Dekkers et al., 2016; Dekkers et al., 2013). First, we trained UDIST on 56,991 PDIOs obtained from a Forskolin Induced Swelling (FIS) assay and treated with different conditions followed by brightfield imaging. The inclusion of varied conditions and multiple imaging timepoints in the FIS assay training dataset ensures broad biological variation, familiarising the model with the range of organoid phenotypes present in these samples. Next, to test the use of UDIST in profiling PDIO phenotypes in relation to therapy treatment, forskolin and DMSO (vehicle control) or CFTR modulators (VX-445/VX-661/VX-770) were added 24 hours prior to brightfield imaging, in line with the protocol described earlier (Lefferts et al., 2024). 1,845 PDIOs were used for UDIST inference, and the extracted latent spaces were all used for downstream analysis. For this dataset, both shape and post-cleanup texture feature spaces were used for downstream analysis, as the two domains are highly correlated in this biological context. Wildtype PDIOs are typically rounder, whereas F508del/F508del PDIOs tend to be more irregular in shape. Notably, even in case where shape features are comparable between conditions, substantial texture differences remain apparent (Figure 5A). Projection of the latent feature spaces revealed a clear separation between conditions (Figure 5B). Inspecting the source images for objects in the PaCMAP shows that the embedding is primarily organised along a size gradient, along which a secondary gradient further separates wildtype from F508del/F508del PDIOs, reflecting the combined contribution of shape and texture to the overall separation of the genotypes and treatment responses. The Sliced Wasserstein Distance quantification shows how the organoids from both wildtype conditions as well as the F508del/F508del + CFTR modulators are nearly indistinguishable from each other in brightfield imaging (Figure 5C). Whereas F508del/F508del + DMSO PDIOs form a uniquely distinct group.

**Figure 5:**
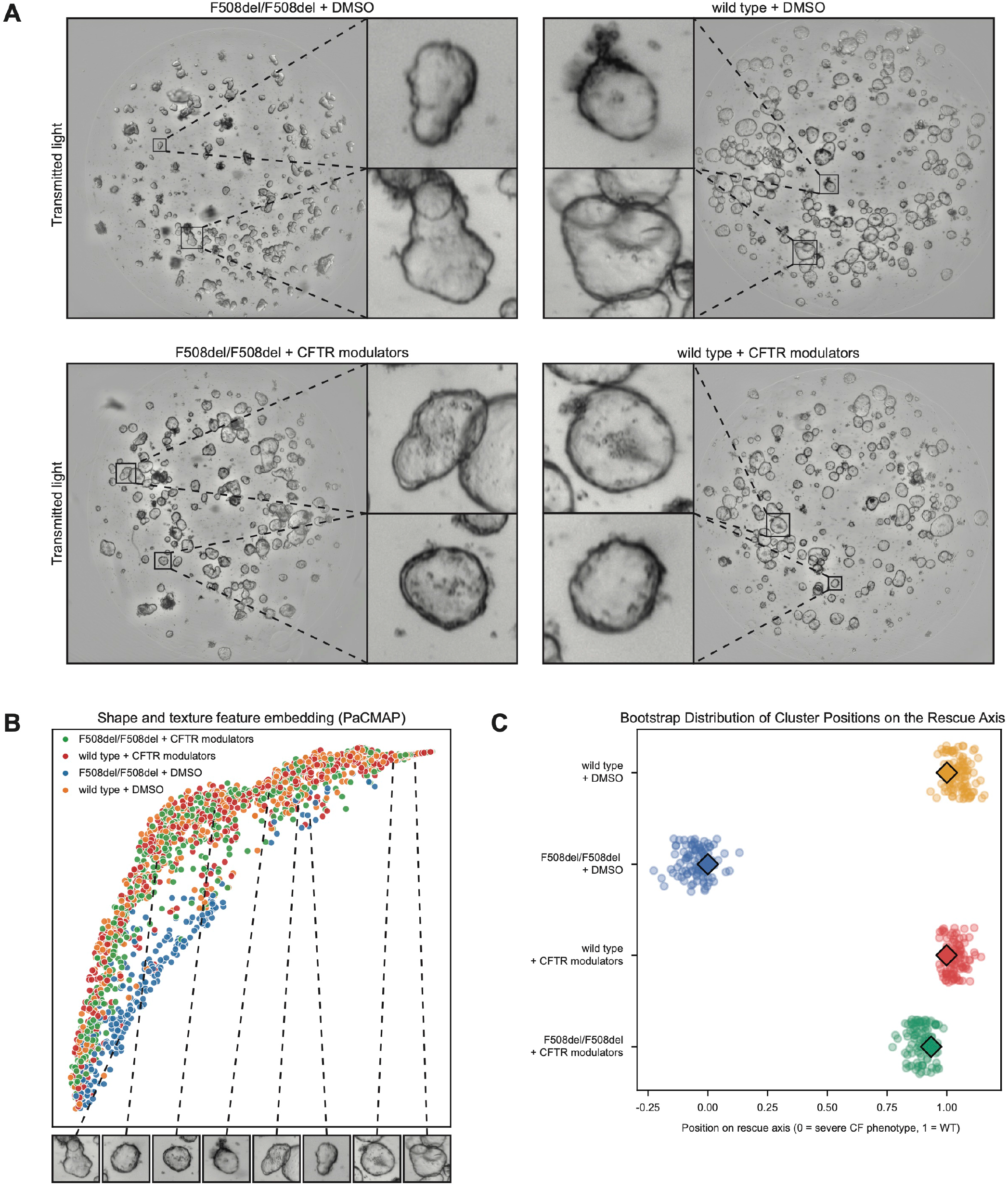
UDIST captures phenotypic differences and treatment-induced rescue in wildtype and F508del/F508del PDIOs from brightfield images. **(A)** Representative brightfield images of wildtype and F508del/F508del PDIOs 48 hours after plating and 24 hours after compound addition. **(B)** PaCMAP embedding of the combined shape and post-cleanup texture spaces, where each point represents a single organoid, coloured by genotype and treatment condition. **(C)** Bootstrap distributions of each condition’s position along the disease–healthy axis, derived from an MDS embedding of baseline-corrected sliced Wasserstein distances between clusters. Diamonds mark the position computed on the full dataset. Points show phenotypic similarity of F508del/F508del PDIOs to wildtype PDIOs under each treatment condition. Scale bars = 30 µm.

Collectively, these results demonstrate that UDIST is not limited to fluorescence imaging but can extract discriminative texture features from brightfield signal. The model successfully distinguishes PDIOs from healthy control donors and F508del/F508del donors, a task in which both shape and texture carry complementary discriminative information. More interestingly, the feature profile of F508del/F508del PDIOs shifts towards that of wildtype PDIOs upon treatment with CFTR modulators, reflecting the known restoration of CFTR function induced by CFTR modulators in F508del/F508del PDIOs (Lefferts et al., 2024)

### Downstream analysis of UDIST latent space using ManiVault Studio

UDIST creates two separate subspaces for each object, shape and texture. To enable users to explore these feature spaces downstream, we recommend using ManiVault Studio (Vieth et al., 2023), a flexible visual analytics tool for analysing high-dimensional data. UDIST allows for the export of shape and texture feature vectors combined with the relevant metadata (e.g. conditions, treatments, cell lines, corresponding images) as a combined CSV file where each row represents a unique object (e.g. nucleus, cell, organoid). This can be loaded into ManiVault Studio (illustrated in Figure S3), where users can analyse the texture, shape, or combined feature spaces, and perform dimensionality reduction accordingly (e.g. t-SNE, UMAP). A particularly powerful feature of ManiVault Studio is its support for interactive drill-down analysis, where users can manually select a subpopulation within an embedding and perform subclustering restricted to that subset, enabling progressively and finer-grained exploration of the latent space. Using the Jupyter Notebook connector of ManiVault Studio, we provide the user with a custom Python notebook for visual inspection of the original images corresponding to a collection of data points within the embedding. Together, this provides users with the means to easily explore the UDIST generated shape and texture latent spaces together with the relevant metadata of their own datasets.

## Discussion

Here we introduced UDIST, a versatile framework for unsupervised, label-free feature extraction in biological image analysis. By creating explicit and independent latent subspaces for shape and texture, UDIST addresses a fundamental limitation of standard AI-based architectures used for autonomous feature extraction in microscopy image analysis: their inability to disentangle these two inherently entangled phenotypic feature domains in an unsupervised manner. This enables researchers to study shape and texture as independent biological readouts.

Existing architectures that conceptually resemble UDIST, such as Conditional Variational Autoencoders (CVAE) (Sohn et al., 2015) and Domain Invariant Variational Autoencoders (DIVA VAE) (Ilse et al., 2020), allow for greater control over feature domain separation, but neither addresses shape-texture disentanglement as a primary objective, resulting in a less explicit separation of shape and texture domains. Furthermore, DIVA VAE requires (semi-)supervised training, limiting its applicability in fully label-free settings. Most closely related to UDIST is MorphoGenie, a recently published unsupervised VAE-based framework for single-cell morphological profiling that achieves disentangled representations across morphological attributes including shape and texture (Murthy et al., 2025). However, MorphoGenie relies on emergent disentanglement, where different latent dimensions incidentally capture different features, rather than explicit architectural separation into dedicated and independent subspaces. Beyond VAE-based disentanglement approaches, UDIST also complements established methods for biological image feature extraction more broadly. Cell Painting CNN is a weakly supervised method designed specifically for Cell Painting datasets that reduces the dominance of shape through contextual cell crops but does not explicitly separate shape from texture features (Bray et al., 2016; Moshkov et al., 2024). O2 VAE achieves rotation-invariant feature extraction but has been shown to retain limitations in texture representation (Burgess et al., 2024), suggesting that UDIST’s explicit texture subspace could serve as a valuable complementary extension. Together, these comparisons highlight that shape-texture disentanglement as an explicit architectural objective remains unaddressed by existing methods. UDIST addresses this directly by guaranteeing shape and texture their own dedicated latent subspaces by design, ensuring complete separation regardless of their degree of correlation in the data.

Using a synthetic dataset with controlled and by design uncorrelated shape and texture features, we demonstrated that shape dominance is a genuine and quantifiable problem in standard VAE-based feature extraction methods. Using UDIST, we showed that explicit shape and texture separation can be achieved by architectural design, with the synthetic dataset serving as a controlled proof-of-concept. This provides a foundation for unsupervised disentanglement of shape and texture in biologically relevant datasets.

To establish biological relevance, we first applied UDIST to a context where texture, rather than shape, contains the primary biological information: the intracellular distribution of EGFR-GFP as a readout for endosomal pathway engagement. We demonstrated that UDIST enables single-channel, label-free endosomal pathway profiling at screening scale. This approach would otherwise be inaccessible to standard feature extraction methods since shape would confound texture features. Importantly, this circumvents the need for exogenous pathway markers such as Dextran and Transferrin, thereby offering a low-complexity and marker-independent approach to endosomal pathway analysis.

We then applied UDIST at the subcellular scale to characterise complex PML-II nuclear condensate phenotypes. As nuclear shape varies between individual nuclei, it risks confounding the analysis of the subtle and heterogeneous PML-II condensate patterns. By studying only PML-II texture patterns, independent of shape, we demonstrated that subtle phenotypic differences within a large screen-based heterogeneous population could be uncovered in an unsupervised manner. This illustrates that disentanglement by design and explicit shape removal are a necessity for the unbiased analysis of subtle phenotypic variation in large heterogeneous screens.

A central assumption underlying UDIST is that shape and texture constitute uncorrelated feature domains. The synthetic dataset was designed to satisfy this assumption explicitly, and the nuclei- and cell-scale biological datasets are broadly consistent with it. The PDIO dataset, however, represents a case in which shape and texture are strongly correlated. Wildtype PDIOs are typically rounder and texturally distinct from CFTR F508del/F508del PDIOs under brightfield imaging. In a standard VAE, such correlation would cause shape to dominate the latent space, leaving texture underrepresented in downstream analysis. UDIST guarantees texture its own dedicated subspace by architectural design, regardless of the degree of correlation with shape. This enabled the detection of a phenotypic shift in F508del/F508del PDIOs towards a wildtype profile upon CFTR modulators treatment: a biologically meaningful result that would likely remain undetected in a standard entangled representation where shape dominates. The organoid dataset therefore demonstrates that UDIST retains its analytical value in biological contexts where shape and texture are correlated and standard methods are likely to underrepresent texture information.

A limitation of the present approach is that cropping to single objects discards the surrounding cellular context, including potential biological information encoded in cell-cell interactions and tissue-level organisation. Per-object analysis also introduces a dependency on the segmentation algorithm, as segmentation errors or inconsistencies will directly propagate into the feature representation. However, a per-object analysis also has an important advantage: by isolating individual objects, the model is better positioned to capture the biological heterogeneity within a population. A further limitation is that UDIST has to date been trained and evaluated exclusively on 2D imaging datasets. Given that biological structures are inherently 3D, extension to volumetric 3D datasets represents a natural and important direction for future development. The current implementation accepts only single-channel images, although the architecture can be readily modified to support multi-channel images.

The present work focused primarily on the UDIST training approach rather than on architecture optimisation; a single general architecture was applied consistently across all datasets. Recent work has proposed evolutionary architecture search as a means of identifying optimal VAE configurations (Shang et al., 2024). A dataset-specific architecture could further enhance UDIST’s performance, although likely at the cost of generalisability. Independently, previous work has shown that loss function design can be of equal or greater importance than model architecture choices (Shan et al., 2023), a finding reflected in the broader VAE literature where most advances have been achieved through modifications to the loss function (Higgins et al., 2016; S. Zhao et al., 2017). To achieve the required disentanglement, UDIST combines multiple loss functions serving distinct objectives: reconstruction quality (masked MS-SSIM, MAE), latent space regularisation (KL-divergence), and representation shaping via VICReg (variance for anti-collapse, invariance for consistency, and covariance reduction for inter-subspace independence). However, as these loss functions pursue competing objectives, residual correlations between shape and texture features are an expected consequence. Recent work exploring multi-objective optimisation strategies to handle competing loss functions in deep learning represents a promising direction to address this (Quinton and Rey, 2024; Nikbakhtsarvestani et al, 2023).

Shape and texture are complementary and biologically informative feature domains for characterising cellular state. Although both are derived from the same underlying object, making their separation inherently challenging, they need not be correlated. Our work demonstrates that through a combination of explicit architectural design and loss function optimisation, VAE models can learn dedicated and independent subspaces for shape and texture, unlike conventional feature extraction methods where both domains remain entangled and their representation quality is consequently compromised. UDIST’s clean separation by design of shape and texture features was first validated on a controlled synthetic dataset with known class structure. Subsequently, this was confirmed across three biological datasets spanning distinct biological scales: nuclear condensates, intracellular receptor trafficking, and patient-derived organoids. These datasets further encompassed two imaging modalities – fluorescence and brightfield – demonstrating that UDIST generalises across diverse high-content microscopy and screening contexts. Collectively, these results establish that shape and texture can be reliably decoupled at the level of individual biological objects regardless of biological scale or imaging modality. Taken together, UDIST provides a versatile and practical tool for unsupervised, label-free microscopy analysis in high-content screening, enabling independent exploration of shape and texture as distinct and biologically informative feature domains at the level of individual objects.

## Methods

### Synthetic data generation

We generate a synthetic fluorescence microscopy dataset consisting of 512*512 centred single-cell-like objects. Each object is assigned a random shape class and a random texture class. We define three shape classes: Round, Elongated and Protrusive. Round cells are generated as closed spline circles with some radial irregularity. Elongated cells are generated as closed spline ellipses with a random aspect ratio (1.8-3.5) between the long and short axis. Protrusive cells are round cells with 2 to 4 radial extensions of variable strength. Each type of cell is generated at a random size scale. All shape types are smoothed by a periodic cubic spline and perturbed by a deformation field to increase edge realism. We define three texture classes: empty, sparse puncta and dense puncta. Each class starts with a base level of random low intensity noise. The sparse puncta class get 3-7 randomly placed high intensity puncta inside the object border based on the mask. For the Dense puncta class we define 3-5 cluster centres at random positions in the cell. At every cluster centre we place between 12 and 20 puncta in a radius of 15 to 30 pixels of the centre only allowing overlap when there are not enough candidate pixels. For every object, regardless of texture class, we apply several filters to increase realism. We apply, subtle vignetting, random illumination gain and offset, random gaussian blur and Poisson-Gaussian noise respectively. We generate 28350 cells for our train set and 8100 cells for our test set

### Endocytosis data

A549 cells (Sigma-Aldrich, CLL1141) expressing EGFR-GFP were seeded on 96 well imaging plates (Revvity, 6055308). On the day of the experiment, cells were serum starved for 2 hours and incubated with the antibody targeting the receptor. Cells were then fixed in 4% PFA (EMS, 15710) containing 1ug/mL Hoechst 33342 (Invitrogen, H3570) for 20 minutes and washed three times in PBS. Images were acquired on a Revvity Opera Phenix Plus spinning disk confocal microscope using a 63x/1.4NA water immersion objective. Three Z planes spaced 1um were collected per FOV and max projected for analysis.

### Nuclei dataset

To generate eGFP-PML-II stable cell lines, plasmids were cloned by Gibson assembly of the PCR amplified eGFP-PML-II inserts into pcDNA5/FRT/TO plasmids. HeLa FlpIn TetR cells were transfected with pOG44 Flp-Recombinase Expression Vector and pcDNA5/FRT/TO eGFP-PML-II plasmid using Lipofectamine 3000 (Thermo Fisher, L3000001). After 2 days, selection was performed using 200 ug/mL Hygromycin. After selection, cells were maintained in 100 ug/mL Hygromycin.

Two custom independent siRNA duplexes were ordered targeting proteins of interest (Sigma Aldrich) and mixed in a 1:1 ratio to prepare a 2uM RNAi pool stock solution in nuclease free water. MISSION® siRNA Universal Negative Control #1 was used (Sigma Aldrich, #SIC001). Transfection mixes were prepared by adding diluted Lipofectamine RNAiMAX (Thermo Fisher Scientific, #13778150) (0.2 μL in 10 μL Optimem) to 1pmol siRNA (0.5 μL of 2 μM stock in 10 μL Optimem) in a 96-well µ-plate (Ibidi, #89606), mixing and incubating for 15 min at room temperature. Subsequently 7000 Hela FlpIn TetR eGFP-PML-II cells were added to each well. Plates were incubated at 37 °C, 5% CO2 for 48 hours after which new medium containing 10 ng/ml Doxycyclin was added to cells to induce eGFP-PML-II expression. After another 24 hours of incubation cells were washed 1x with PBS and fixed with 4% PFA for 10 minutes at room temperature. After fixations cells were washed with PBS and permeabilized with 0.2% Triton X100 in PBS for 10 minutes at room temperature. Cells were subsequently washed with PBS and incubated for 30 minutes at room temperature with 2% BSA in PBS (blocking buffer). Cells were incubated with primary antibodies in blocking buffer for 2 hours at RT. The following primary antibodies were used: mouse anti-lamin B1 (Santa Cruz, sc-365214, 1/500) and mouse anti-lamin A/C (Santa Cruz, sc-376248, 1/500). Cells were washed 3x with PBS and incubated with Alexa Fluor 647 goat anti-mouse IgG2b (Life technologies, #A21242), Alexa Fluor 568 goat anti-mouse IgG1 (Life technologies, #A21124) and DAPI for 1 hr at room temperature. Cells were washed 3x with PBS, overlaid with 100 μL/well PBS and stored at 4 °C until imaging. The experiment was performed in triplicate.

Imaging was performed on an Operetta CLS High-Content Analysis System (Revvity) using a Andor Zyla 5.5 camera and a 63x water immersion objective (NA 1.15) in spinning disk mode. The following channels were recorded: DAPI (Excitation 355 – 385 nm, Emission: 430 – 500 nm), EGFP (Ex: 460 – 490 nm, Em: 500 – 550 nm), Alexa 568 narrow (Ex: 530 – 560 nm, Em: 585 – 610 nm) and Alexa 647 (Ex: 615 – 645 nm, Em: 655-760 nm). 5 z-slices were taken 0.7 µm apart and max projected for analysis.

### Cystic Fibrosis PDIO dataset

#### Forskolin induced swelling (FIS) assay

Organoids were cultured in 50% Matrigel droplets and passaged weekly using mechanical disruption, as described previously (Lefferts et al., 2024). Cultures were maintained in medium supplemented with Wnt3a alternative peptide (peptigrowth, PG-008). Organoids were manually disrupted prior to plating. Subsequent steps of the assay were performed using an automated pipeline. Disrupted organoids were plated into 96-wells plates using a liquid handler (Opentrons OT-2). After plating, standard intestinal culture medium was added using a second liquid handler (Biotek Multiflo). After a 24-hour recovery period organoids were treated with CFTR correctors: VX-121 (SelleckChem, Cat# E1701), VX-445 (MedChemExpress, Cat# HY-111772/CS0090942), VX-661 (MedChemExpress, Cat# HY-15448) or VX-809 (MedChemExpress, Cat# HY-13262). After an additional 24 hours, CFTR potentiators, VX-770 (MedChemExpress, Cat# HY-13017/CS-0497) and VX-561 (SelleckChem, Cat# E1743) were added, together with forskolin (Sigma Aldrich, Cat# F3917). All CFTR modulators were used at a final concentration of 3 µM. The final concentrations forskolin were 0.02, 0.128 and 0.8 µM, with DMSO included as a vehicle control. Compounds were dispensed into the 96-wells plates using a liquid handler (Dispendix, I.DOT). Immediately after adding forskolin, Forskolin induced swelling was tracked for one hour, with repeated acquired images every ten minutes using a Zeiss Cell Discoverer 7 (CD7) microscope at 2.5x magnification.

#### Drug induced swelling (DIS) assay

Organoids were cultured in 50% Matrigel droplets and passaged weekly using mechanical disruption, as described previously (Lefferts et al., 2024). Cultures were maintained in medium supplemented with Wnt3a alternative peptide (peptigrowth, PG-008). Organoids were manually disrupted prior to plating. Subsequent steps of the assay were performed using an automated pipeline. Disrupted organoids were plated into 96-wells plates using a liquid handler (Opentrons OT-2). After plating, standard intestinal culture medium was added using a second liquid handler (Biotek Multiflo). After a 24-hour recovery period organoids were treated with forskolin (Sigma Aldrich, Cat# F3917) and the CFTR modulators: VX-770 (MedChemExpress, Cat# HY-13017/CS-0497), VX-445 (MedChemExpress, Cat# HY-111772/CS0090942), and VX-661 (MedChemExpress, Cat# HY-15448). A final concentration of 3 µM was used for all CFTR modulators, and 0.8 µM of Forskolin was used along with a DMSO control. Compounds were dispensed into the 96-wells plates using a liquid handler (Dispendix, I.DOT). Brightfield images were acquired 24 hours after treatment using a Zeiss Cell Discoverer 7 (CD7) microscope at 2.5x magnification.

#### Ethical approval

Ethical approval for use of organoids: Organoids used were obtained from the HUB (Hubrecht Organoid Technology) Biobank (www.huborganoids.nl) under TC-Bio protocol number 14-008, or from the UMCU Darmbank under TC-Bio protocol number 19-831, and used according to informed consent.

### Image preprocessing

For every dataset both images with objects and labelled masks are required. The synthetic dataset was generated from masks, so no segmentation was needed. The endocytosis dataset and nuclei dataset were segmented using Cellpose 3.0.6 on default settings using the nuclei channel and the channel of interest (Stringer et al., 2021). Segmentation of the organoid dataset was performed using a retrained Cellpose 4.0.2 (Pachitariu et al., 2025) on the dataset introduced by Lefferts et al (Lefferts et al., 2024). Using the masks, the objects were cut out in the channel of interest. The object crop and the mask crop were then zero-padded to a set size of 512*512 pixels. After cropping each object-mask combination was aligned to the principal axis as implemented by (Pedregosa et al., 2011).

### Architecture

The basic VAE model used is a convolutional neural network (CNN) based architecture consisting of encoder and decoder blocks. The model starts by performing a convolution step with a kernel of size (7,7) and 64 output channels. Followed by four residual blocks with output channels 64,128, 256, 512. Each of the residual blocks performs convolution, batch norm, ReLu, convolution, batch norm, ReLu. Each residual block additionally implements a skip connection following the principles described in the resnet paper (He et al., 2016). The final 512 channel 32*32 images are flattened and passed through the model bottleneck. The VAE bottle neck consists of two parallel linear layers to project the input to a mean and a logvar vector. These are reparametrized following the standard approach from the original VAE implementation (Kingma & Welling, 2013). The decoder blocks are constructed in such a way as to avoid any checkboard artifacts (Odena et al., 2016). Each block performs convolution, batch-norm, ReLu, upsample, convolution, batch-norm, ReLu. The upsample layer uses nearest neighbour upsampling for shape models and bilinear upsampling for texture models.

### Loss function

The loss function is composed of 6 metrics calculated at various points in the model. In no particular order, the metrics are : Variance-Invariance-Covariance regularisation (VICReg) (Bardes et al., 2021), L1 loss, Masked MS-SSIM (Wang et al., 2003) and KL divergence. Variance, Invariance, Covariance and the Kullback-Leibler divergence each have their own weight determined through hyperparameter tuning. Namely, 10 (lambda_var), 100 (lambda_inv), 0.1 (lambda_cov) and 0.01 (lambda_kl) respectively. L1 and Masked MS-SSIM are combined to form a single reconstruction loss as defined by

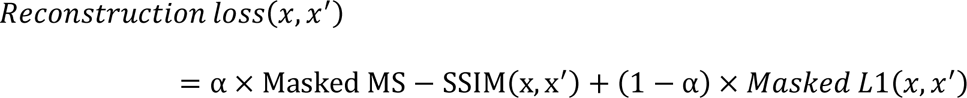

Where α is set at 0.84. This specific loss is based on (H. Zhao et al., 2017). Due to the centred, masked and padded nature of our images we adapt both the MS-SSIM loss and L1 loss to be masked implementations. This combination of L1 and MS-SSIM is weighed together as the reconstruction loss in the final formulation with a value of 10000 (lambda_recon). We calculate the Variance, Invariance and Covariance between latent vectors of the original and an augmented image as described by Bardes et al. (Bardes et al., 2021). The Kullback Leiber divergence was calculated as defined by the original VAE paper. (Kingma & Welling, 2013)

### Model training

Model training is performed in a two-step process. The first step is training a full VAE on the zero padded cutout binary masks for each individual object in the training set. The VAE is trained using the loss function as described before to achieve descriptive structured latent vectors. This model is trained until the training loss functions stabilize, at which point the weights are frozen. The second step consists of training a second identical VAE on the zero padded cutouts of the actual objects. During training of this second VAE, for every object we also encode the corresponding mask. The new decoder is fed both the latent vectors produced by both the new and the frozen encoder.

### Hyperparameter tuning

Because each of the loss components attempts to optimize separate parts of the training and often end up competing, we perform hyperparameter tuning as a multi-objective optimization problem. Using Optuna, an open source hyperparameter optimization framework, we searched for the pareto frontier (Akiba et al., 2019). We evaluated the following components: variance weight, covariance weight, invariance weight, KL weight, reconstruction weight and latent space size. This eventually let us define a one-size-fits-all set of loss weights and latent space size that performed well for each of our datasets.

### Residual information subtraction

We remove any residual shape information from the intensity latent vectors by Principal Component Regression. This method first performs Principal Component Analysis (PCA) on the full test set of shape latent vectors generating as many principal components as there are shape latent dimensions. Then the Principal Components are regressed against the intensity latent vectors using Ordinary Least Squares regression. This results in the component of the intensity based variation that can be linearly explained by the shape components. We then subsequently subtract this predicted component from the intensity latent vectors to clean any shape related leakage during model training.

### PaCMAP

Dimensionality reduction is performed as described and implemented by Wang (Wang et al., 2021). For every plot we used the following settings: 50 nearest neighbours (n_neighbors=50) with the ratio of mid near pairs to nearest neighbour pairs set at 0.1 (MN_ratio=0.1) and the ratio of further pairs to nearest neighbour pairs at 2.0 (FP_ratio=2). The distance metric was angular (distance=’angular’) with the reducer being initialised with Principal Component Analysis (init=’pca’)

### Sliced Wasserstein Distance

SWD treats every cluster as a probability distribution and calculates the similarity between these clusters. By projecting the data onto random one dimensional axes and averaging the one-dimensional Wasserstein distance over all axes it calculates a distance score. We normalize the results between 0-100 to ensure valid comparisons between experiments. We calculate the SWD as follows:

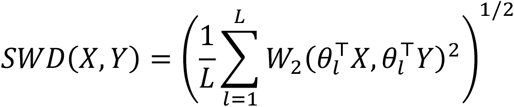

Where L is a hyperparameter for the number of projections to sample, θ*_l_^T^* is a 1D projection of the high dimensional space in a random direction and W_2_ is the 1D Wasserstein distance. Additionally we implement a bias subtraction to account for cluster noise. We split each cluster in our data in two random halves over which we calculate the SWD. We do this 20 times per cluster. We then calculate the final distance score as

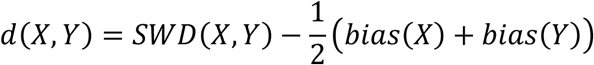

### Leiden Clustering

To obtain a neighbourhood graph from our latent vectors we use UMAP with 128 neighbours to create a fuzzy simplicial complex graph. We ran the Leiden algorithm on this graph with a default resolution of 1. To account for the stochastic nature of the Leiden algorithm we perform 5 independent runs with different seeds. A final consensus graph was derived from these runs by only keeping edges that cluster together over 50 percent of the time (at least 3 out of 5 runs). We then run Leiden clustering a final time over this consensus graph to obtain our final labels.

### Leiden correspondence significance test

For each combination of Leiden cluster and siRNA duplex, a log₂ enrichment score was calculated as the log₂ ratio of the observed frequency of a given duplex’s nuclei within a Leiden cluster to the expected frequency of nuclei in that cluster, where the expected frequency was defined by the distribution of siCTRL nuclei across Leiden clusters. A chi-square test of independence was performed to assess the significance of each cluster-duplex pair, except where the minimum observed count in the corresponding 2×2 contingency table was below 5, in which case a Fisher’s exact test was performed instead. The Benjamini-Hochberg false discovery rate (FDR) procedure (Benjamini & Hochberg, 1995) was used to correct for multiple comparisons across the resulting p-values, with a significance threshold of α = 0.05.

### Multidimensional scaling to a rescue axis

To generate a rescue axis a bootstrap procedure was performed (100 iterations). In Each iteration 80% of each cluster’s points were sampled. The Sliced Wasserstein Distance was calculated over this subset as explained before. The resulting distances were embedded in two dimensions using metric Multidimensional Scaling (Borg & Groenen, 2005; Torgerson, 1952) and aligned by rotation and reflection using Kabsch algorithm without rescaling (Kabsch, 1976). We then defined a one-dimensional rescue axis from the reference position of DMSO F508del/F508del to reference position of DMSO WT and projected all points on that axis.

## Data availability

The training and experimental data that support the findings of this study are available upon request during peer review and will be made publicly available on Zenodo (https://zenodo.org/communities/livingtech/) upon publication.

## Code availability

The optimized code is available under the GPL-3 license at: https://github.com/Living-Technologies/UDIST

The code includes an example config file with a corresponding reader and label extractor for the synthetic dataset.

## Acknowledgements

This work is part of a Health∼Holland sponsored public-private partnership between UMC Utrecht, Genmab B.V. (providing additional in-cash sponsoring), and Utrecht University. This work was further financially supported by the strategic alliance between Eindhoven University of Technology (TU/e), Wageningen University & Research (WUR), Utrecht University (UU), and UMC Utrecht, and by the Dutch Research Council (NWO) as part of the project Innovative Stem Cell Technology Infrastructure for Human Organ and Disease Models (project number 184.036.006) of the Large-Scale Research Infrastructure programme. A.F.J.J. was supported by a Leverhulme Trust Early Career Fellowship and the Isaac Newton Trust. We thank Heather Zecchini from the Light Microscopy Core Facility (RRID:SCR_028019) at the CRUK Cambridge Institute for expert assistance with imaging.

## Author contributions

B.M.B. designed the study, coordinated the project, designed and developed the method, wrote the code base, analysed the results, and wrote the manuscript; M.L.T. co-designed and co-developed the method, provided AI expertise, and supervised the research; M.B.S. co-designed and co-developed the method, provided image analysis expertise, and reviewed the method and code base; K.H.v.d.S. provided interactive tooling for downstream analysis; C.T.H.J. provided high-content microscopy expertise, designed and performed EGFR-GFP related (microscopy) experiments, and reviewed the method; V.O. and T.H.W. performed EGFR-GFP related (microscopy) experiments; A.F.J.J. designed and performed PML-II related (microscopy) experiments and reviewed the method; L.W. performed organoid related cultures and (microscopy) experiments; E.M.A.H. reviewed the method and code base; J.W.L., E.M., and E.D.E. provided high-content microscopy expertise and reviewed the method; C.A.T.v.d.B. co-designed and co-developed the method, supervised the research, and provided AI expertise; J.M.B. co-designed and co-developed the method, supervised the research, and provided CF and organoid expertise; S.F.B.v.B. co-designed and co-developed the method, provided AI and microscopy expertise, supervised the research, and wrote the manuscript. All authors reviewed the manuscript.

## Declaration of generative AI and AI-assisted technologies in the writing process

During the preparation of this work, the authors used Claude (Anthropic) for copy editing to improve language and readability. After using this tool, the authors reviewed and edited the content as needed and take full responsibility for the content of the publication.

---

**Supplementary Figure 1:**
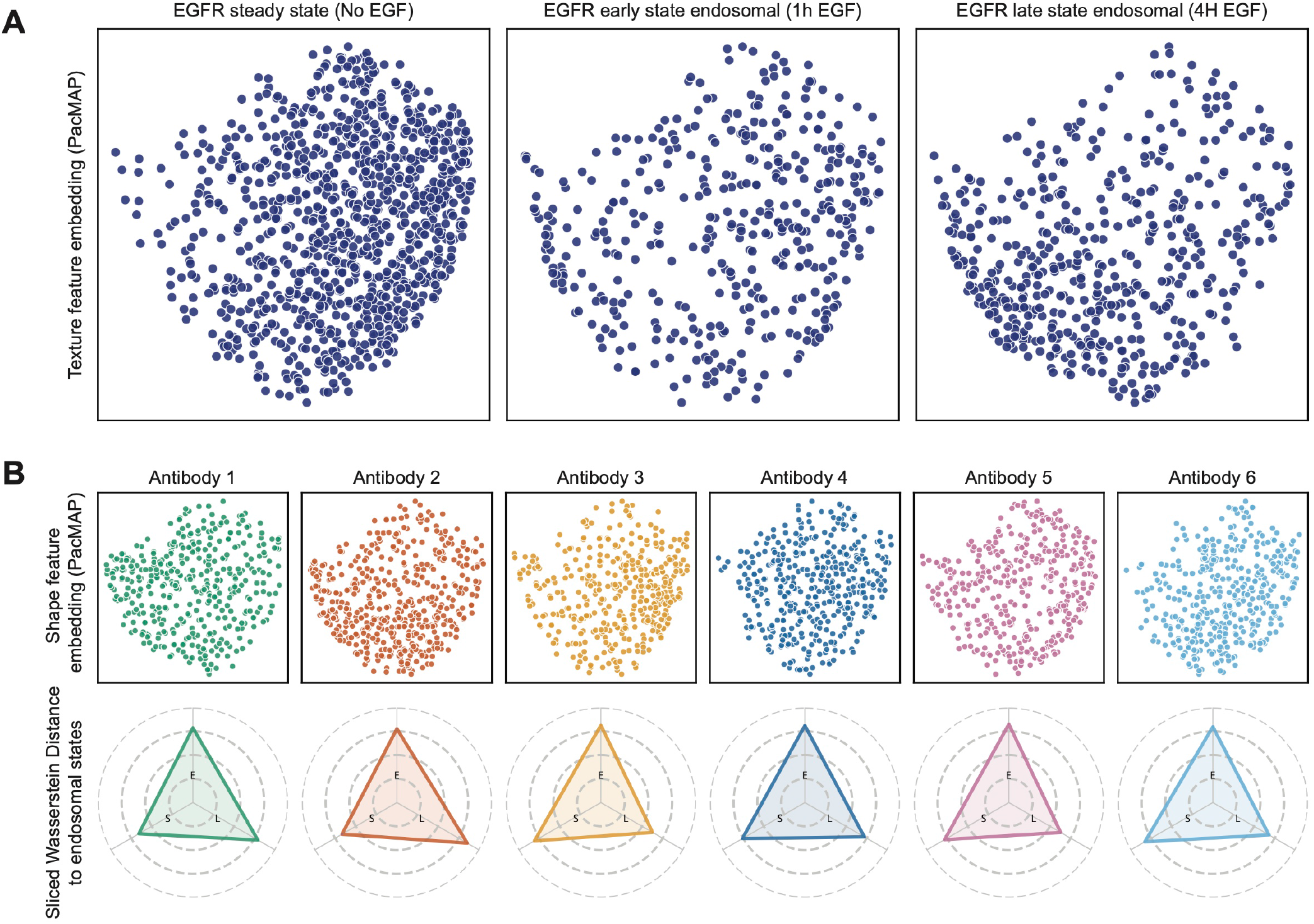
Shape features show a uniform distribution across EGFR-GFP experimental conditions. **(A)** PaCMAP embedding of shape latent features for all cells in the control conditions, where each point represents a single cell. **(B)** PaCMAP embedding of the shape latent features for each experimental condition, where each point represents a single cell (upper panel). Similarity scores of each experimental condition relative to each control condition, calculated in the original latent space using the Sliced Wasserstein Distance and min-max normalised to the measured texture distances (lower panel). S: steady state, E: early endosomal, L: late endosomal. PaCMAP embeddings show a random subsample of 10% of all cells.

**Supplementary Figure 2:**
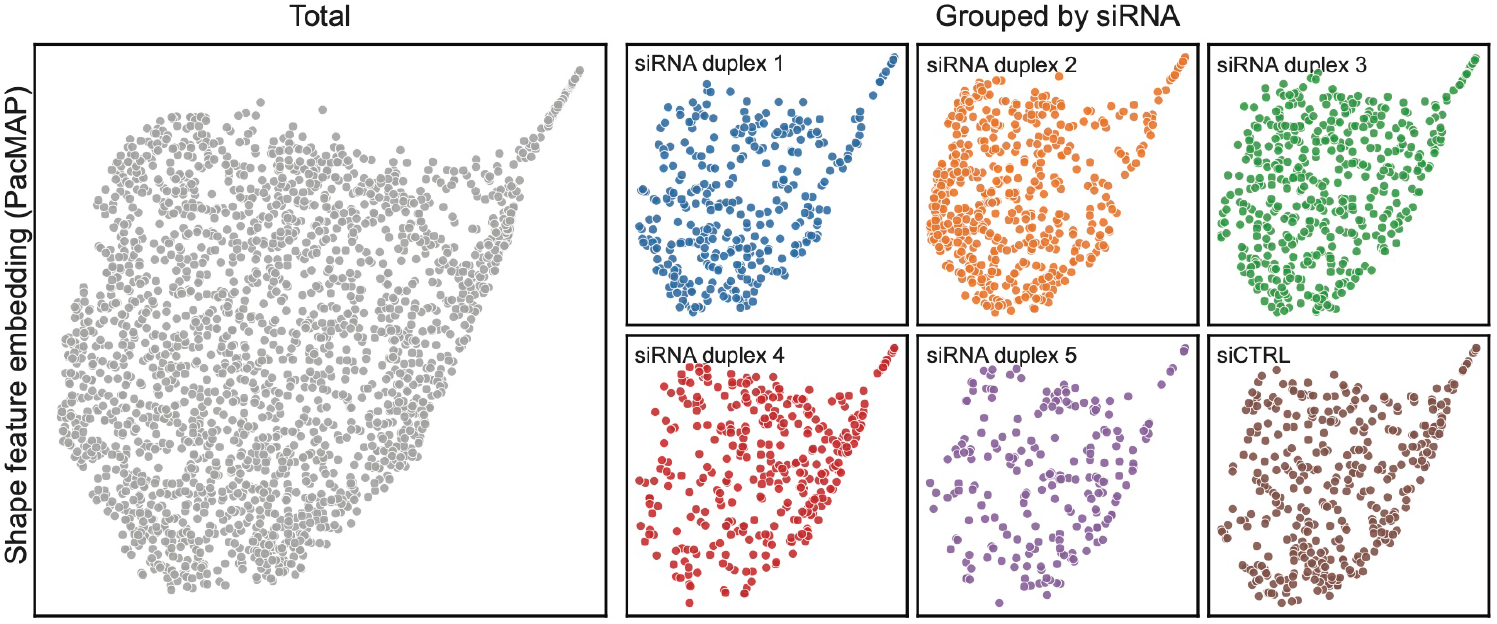
Shape features show a uniform distribution across GFP-PML-II siRNA conditions. PaCMAP embedding of shape latent features where each point represents a single nucleus (left panel), and PaCMAP embeddings showing localisation of each siRNA condition within the shape latent space (right panels).

**Supplementary Figure 3:**
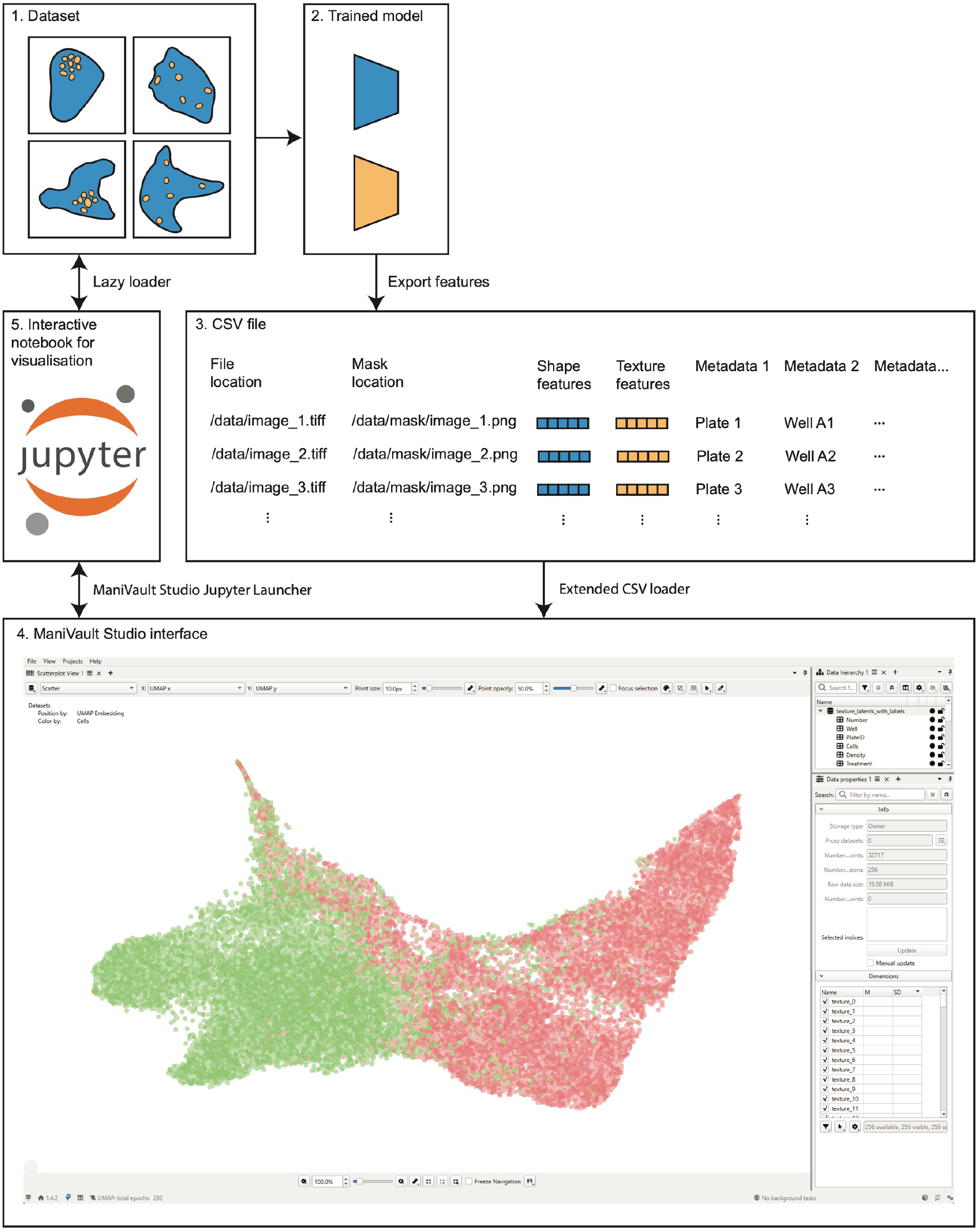
Schematic overview of the ManiVault Studio workflow for downstream analysis of UDIST latent spaces. Schematic overview of the recommended workflow for downstream analysis of UDIST latent vectors using ManiVault Studio. (1) The dataset is stored on a cloud-based server, local server or local device. (2) A UDIST model is trained on the data and the model weights are saved. (3) Extracted feature vectors, relative file paths to the corresponding original images, and associated metadata are stored in a combined CSV file. (4) The CSV file is loaded into ManiVault Studio for interactive exploration and analysis of the latent spaces. (5) A Jupyter notebook connector can be opened for visual inspection of original images corresponding to selected data points within the embedding.

